# ESPRESSO: Spatiotemporal omics based on organelle phenotyping

**DOI:** 10.1101/2024.06.13.598932

**Authors:** Lorenzo Scipioni, Giulia Tedeschi, Mariana Navarro, Yunlong Jia, Scott Atwood, Jennifer A. Prescher, Michelle Digman

## Abstract

Omics technologies, including genomics, transcriptomics, proteomics, and metabolomics, have been instrumental to improving our understanding of complex biological systems. Despite fast-pace advancements, a crucial dimension is still left to explore: time. To capture this key parameter, we introduce ESPRESSO (Environmental Sensor Phenotyping RElayed by Subcellular Structures and Organelles), a pioneering technique that provides high-dimensional phenotyping resolved in space and time. Through a novel paradigm, ESPRESSO combines fluorescent labeling, advanced microscopy and bioimage and data analysis to extract morphological and functional information of the organelle network unveiling phenotypic changes over time at the single-cell level. In this work, we present ESPRESSO’s methodology and its application across numerous cellular systems, showcasing its ability to discern cell types, stress response, differentiation and immune cells polarization. We then correlate ESPRESSO phenotypic changes with gene expression and demonstrate its applicability to 3D cultures, offering a path to revolutionizing biological exploration, providing invaluable insights into cellular states in both space and time.

## Introduction

We live in the midst of a historical moment, witnessing omics technologies (particularly transcriptomics^1^, proteomics^2^, and metabolomics^3^) transitioning from homogenized “bulk” measurements to the intricacies of single-cell and spatial analyses. While informative, bulk approaches mask the inherent heterogeneity within cellular subpopulations. Single-cell techniques like single-cell RNA sequencing (scRNA-seq) have overcome this limitation, unveiling distinct cell types, rare subpopulations, and insights into complex biological processes. However, scRNA-seq lacks spatial information, hindering our understanding of how individual cells organize and interact based on their relative positions. Spatial omics^4^ represents the natural evolution of these technologies, revealing how phenotypes are spatially organized as well as communication networks between cells. Despite these advancements, capturing the dynamic aspects of these interactions remains a significant challenge. Indeed, transitions between observed phenotypes are just as important, if not more important, than their mere presence. Numerous cellular processes leading to phenotypic changes occur over time, regulating development (e.g., differentiation), immune response (e.g., macrophage polarization), or adaptation to external conditions (e.g., stress response and metabolic switch). While the pursuit of spatiotemporal omics holds immense promise, adapting traditional approaches and achieving true spatiotemporal resolution presents significant challenges that must be addressed in order to achieve true understanding of these biological processes.

Current omics techniques, while informative, often fall short in capturing the dynamic nature of biological transitions, particularly in metabolic shifts where cells undergo rapid reprogramming of gene expression, protein activity, and metabolic pathways. To unravel these complexities and identify potential therapeutic targets, time-resolved omics data with high-temporal-resolution is indispensable, rendering imperative to develop faster, more sensitive spatial omics techniques.

Here, we propose a paradigm shift towards a different phenotyping target: organelles. Extensive literature links organelle properties to specific cell states^5^. As an example, organelle characteristics are tightly linked to cellular metabolic processes^6^ and differentiation state^7^ and can inform on the cell’s bioenergetics^8^. They can identify the cellular response to specific stressors (metabolic stress^9^, mitochondrial fragmentation^10^, mitophagy^11^, lipid storage^12^ and consumption^13^), while lipid droplet characteristics (e.g., number and size) have been linked to proliferation and quiescence in neural stem cells^14^ and to specific breast cancer stem cells phenotypes^15^. In a fascinating process called organelle inheritance^16^, daughter cells with different fates (e.g., differentiation state) inherit organelles with different characteristics, hinting at the organelle network’s importance in maintaining cellular homeostasis. Techniques like the Cell Painting^17^ assay leverage organelle properties using fluorescent dyes to target various components like the nucleus, endoplasmic reticulum, mitochondria, cytoskeleton, Golgi apparatus, and RNA, and demonstrate applications in drug discovery. Proteomics has also been applied in organelle-specific contexts with techniques like nuclear proteomics^18^, where nuclear isolation followed by mass spectrometry analysis unveils protein composition and interactions within the nucleus. Overall, a clear link exists between the morphological and functional characteristics of the organelle network (hence, the “organelle landscape”) and cell state.

Here, we developed an innovative approach called ESPRESSO (Environmental Sensor Phenotyping RElayed by Subcellular Structures and Organelles), providing high-dimensional phenotyping resolved in space and time, thus overcoming the above limitations. To this aim, we assembled a panel of four organelle-specific, live-cell dyes with signals stable for at least 24 hours. Furthermore, we selected dyes that modify their fluorescence intensity based on the biophysical properties of their target organelle (e.g., lysosomal acidity^19^ and mitochondrial membrane potential^20^), simultaneously reporting on both morphological (e.g., number, size, interaction and subcellular organization) and functional information from four organelles simultaneously. We imaged the labeled cells using hyperspectral imaging^21^, a fluorescence microscopy technique that acquires the entire fluorescence emission spectrum pixel-by-pixel, allowing snapshot multicolor imaging. We then proceeded to combine this labeling and imaging strategy with convolutional neural networks, signal unmixing, bioimage analysis, and data analysis tools, thus creating the ESPRESSO framework. In the following sections, we present ESPRESSO’s methodology and its application across various cellular systems, showcasing its ability to discern different cell types, phenotypic adaptation to stressor exposure, keratinocyte differentiation, and macrophage polarization. We then demonstrate that ESPRESSO’s phenotypic changes correlate with gene expression and feature its applicability to 3D triple-negative breast cancer spheroids, analyzing their phenotypic evolution during collagen invasion and response to various stressors. ESPRESSO represents a framework that offers high-dimensional phenotyping based on organelles, representing the first spatiotemporally-resolved omics technique based on this approach.

## Results

### ESPRESSO workflow and cell type phenotyping

Development of a comprehensive method for spatiotemporal omics requires multiple techniques to merge into a unified framework and is schematized in Figure 1a.

**Figure 1.**
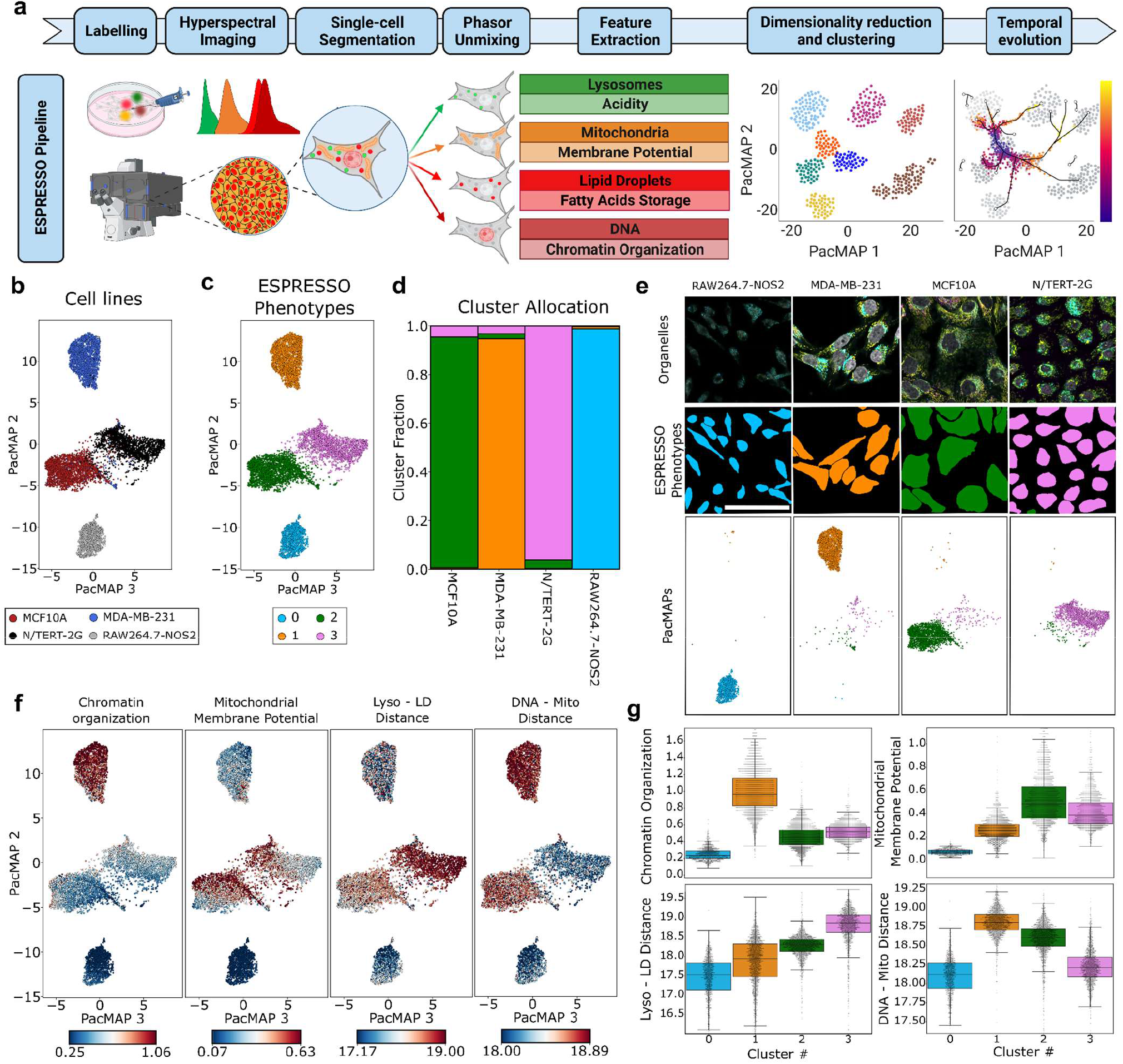
ESPRESSO workflow and organelle phenotypes of diverse cell lines. Schematic representation of the ESPRESSO workflow (a). Cells are labeled with a panel of live-cell fluorescent dyes targeting lysosomes, mitochondria, lipid droplets and DNA. Cells are then imaged with confocal microscope equipped with hyperspectral detection and single cells are isolated from images by segmentation. The images relative to individual organelles are unmixed by spectral phasor unmixing and morphological and functional properties of the organelle network are extracted, dimensionally reduced and clustered to identify organelle phenotypes. Subsequently, the temporal evolution of ESPRESSO phenotypes is monitored over time. ESPRESSO PacMAP of 8,838 cells from four different cell lines (MCF10A, MDA-MB-231, N/TERT-2G and RAW264.7-NOS2) color-coded by cell line (b) and ESPRESSO phenotypes (c). Stacked bar plot showing the cluster fraction for each ESPRESSO phenotype (d). Representative regions of interest (e) showing the unmixed organelles images (top, white: DNA, yellow: mitochondria, cyan: lysosomes, magenta: lipid droplets), the cell masks color-coded by ESPRESSO phenotype (center) and corresponding ESPRESSO PacMAPs (bottom). ESPRESSO PacMAPs color-coded by representative organelle properties (f), calculated as the center of mass of the intensity distribution, cross- or auto-correlation curves, as described in the Methods section. Swarm plots and box-and-whiskers plots (g) quantifying the properties shown in f for each cluster. The black central line in the box-and-whiskers plot denotes the median value, the box extends between the upper and lower quartile and the whiskers extend between the minimum and maximum value, excluding outliers. Scalebars in e are 50 µm. LD: Lipid Droplets, Mito: Mitochondria, Lyso: Lysosomes

#### Organelle-specific, environment-sensitive dyes

First, we worked on defining a panel of four organelle-targeting, live-cell compatible fluorescent probes and optimized the staining protocol to achieve >24-hour continuous imaging with stable fluorescence signal and spectral signature in living cells (Figure S1). The emission spectra of the dyes panel, however, were highly overlapping (Figure S1) and precluded signal discrimination with conventional multicolor microscopy techniques (e.g. multiple excitations and emission filter sets), that have the further disadvantage of performing multicolor imaging sequentially, which would hinder the quantification of the relative position of different organelles within the cells.

#### Hyperspectral imaging and phasor unmixing

To overcome this limitation, we turned to hyperspectral imaging^22^, a technique based on confocal microscopy that allows for the simultaneous acquisition of the emission spectrum across the visible range in a pixel-wise manner. Once the hyperspectral image is acquired, we applied spectral phasor unmixing^23,24^ to separate the signal of the individual probes, allowing for the simultaneous imaging of the four organelle-targeting fluorophores in our panel, regardless of spectral or spatial overlapping. At this stage, we confronted the challenge presented by the imaging speed. Indeed, the higher the number of photons, the better the unmixing and quantification of the dyes’ response, but also the higher is the phototoxicity to the sample.

#### Neural networks

To solve this conundrum, we turned to convolutional neural networks (CNNs), now ubiquitous in image analysis^25^ thanks to their high performance and versatility. We leveraged that power by training a content-aware image restoration (CARE^26^) denoising convolutional neural network to increase the acquisition speed 16-fold while maintaining image quality (Figure S2). Furthermore, we introduced a cell-segmentation CNN (CellPose^27^) in our pipeline to achieve single-cell capability via cell segmentation.

#### Single-cell features extraction

Once segmented, we addressed the extraction of the organelle features from each of the single cells. After unmixing, the images of the target organelles (lysosomes, mitochondria, lipid droplets and DNA) were analyzed morphologically by image correlation spectroscopy (ICS^28,29^), while intracellular organelle distribution was assessed by image cross-correlation spectroscopy (ICCS^30,31^). ICS and ICCS are well-established methods to investigate the number and size of subcellular structures^32–34^ and their colocalization or distance within the cell. Despite all information about these important morphological features is indeed embedded in the ICS and ICCS functions, a model is traditionally applied to quantify them, forcing us to make assumptions about the organelles under analysis. Aiming to construct ESPRESSO as a general framework, we by-passed modeling and fitting and we leveraged the entire ICS and ICCS functions, considering each bin of their rotational average as a morphological feature. A similar approach was taken to extract functional information. Indeed, each of the selected organelle dyes’ intensity is dependent on a defined property of the target organelle (acidity for lysosomes, membrane potential for mitochondria, fatty acids storage for lipid droplets and chromatin organization for DNA). Similarly to morphological features, we assessed functional features by computing the intensity distribution of each unmixed organelle image, therefore taking into account not only mere descriptors of the distribution, but the distribution in its entirety, that encodes information about the heterogeneity in the function of individual organelles, as described in detail in the Methods section. As a result, a collection of 3,328 features was obtained reporting on morphology, intracellular organization and function, that can be carried over to the final step in our pipeline, which concerns the analysis of the organelle profiles.

#### Data analysis and clustering

Similarly to gene expression profiles in scRNA-seq, organelle properties are normalized, selected and reduced in dimensionality by PacMAP^35^, generating low-dimensional embeddings that encode the high-dimensional organelle properties of each cell. A Gaussian Mixture Model (GMM^36^) clustering algorithm is then applied and the number of clusters is based on the minimization of the Davies-Bouldin Index^37^, as described in the Methods (Figure S3, S4). We will refer to these clusters, that identify cell states based on organelle properties, as ESPRESSO phenotypes. In conclusion, thanks to a combination of stable organelle-targeting dyes, simultaneous multicolor imaging and denoising, single-cell segmentation, bioimage analysis and data analysis tools, we obtained the ESPRESSO phenotypes at the single-cell level and can assess their evolution as a function of time for over 24 hours (for a detailed workflow, see Figure S5), as we demonstrate by discriminating between different cell types (Figure 1).

#### Proof-of-concept experiment

In this experiment, we analyzed four different cell lines through the ESPRESSO pipeline previously described. We plated MDA-MB-231 (triple-negative breast cancer), MCF10A (non-tumorigenic epithelial), N/TERT-2G^38^ (immortalized keratinocytes) and RAW264.7-NOS2^39^ (mouse macrophages expressing a NOS2 gene expression reporter) cells in growth conditions and segmented 8,838 cells for ESPRESSO phenotyping. As shown in Figures 1c and 1d, GMM clustering easily identified the cell type-specific phenotypes and allowed the quantification of properties of interest in their organelle network. As an example, we can appreciate from the heatmaps in Figure 1e that MDA-MB-231 cells are characterized by a higher chromatin compaction when compared to the other cells lines, that are instead characterized by high mitochondrial membrane potential (MCF10A), high lysosome-lipid droplet distance (N/TERT-2G) or low chromatin organization and low mitochondrial membrane potential (RAW264.7-NOS2). Finally, ESPRESSO phenotyping, being based on microscopy imaging, is intrinsically spatially-resolved and cells can be color-coded based on the assigned ESPRESSO phenotype (Figure 1f) and, since the information is obtained in high throughput from living cells, the changes in their phenotypes can be followed and analyzed over extended periods of time.

### ESPRESSO phenotypic response to stressor exposure

After establishing that ESPRESSO can discriminate between cell types, we moved on to studying the effect of stressor exposure. Stress response is the cellular response to changes in environmental conditions and involves a multifaceted reprogramming of cellular processes to adapt to and overcome new environments. As an example, hypoxia (oxygen deprivation) reduces the cell’s ability to generate energy through aerobic respiration^40^, leading to a signaling cascade that ultimately results in a metabolic reprogramming towards a more anaerobic energy production. Understanding how different cell lines behave in response to different stressors can provide valuable insights into cellular resilience and has potential for therapeutic interventions aimed at modulating stress responses in disease states. Here, we exposed MCF10A cells to a number of chemicals that changed the environmental conditions towards hypoxia (CoCl_241_), cholesterol inhibition (simvastatin^42^), autophagy inhibition (bafilomycin A1^43^), mitochondrial stress (FCCP^44^) and chromatin compaction (anacardic acid^45^) or decompaction (trichostatin A^46^) for 24 hours and measuring the resulting ESPRESSO phenotype. Our aim is to demonstrate that the wealth of information obtained by ESPRESSO phenotyping can not only detect stress responses, but can also distinguish between different types of stress. Indeed, we expect the different treatments to activate distinct cellular stress response pathways, leading to distinct phenotypes, as a result of the environmental changes we introduced. From the ESPRESSO PacMAPs (Figure 2a and b), we can identify a major group of cells (center) and two distinct smaller groups that we can associate with cells containing large lysosomes (bottom) and with cells displaying compacted chromatin (top), as can be appreciated in Figure 2d and 2e, speaking to the intrinsic heterogeneity of the cell population in itself. As for the ESPRESSO phenotypic response to our panel of stressors, we observed two high-order effects across these chemicals: they either altered the allocation of phenotypes that also exist in control conditions (CoCl_2_ and Simvastatin) or they introduced new, stress-specific phenotypes (e.g., trichostatin A, bafilomycin A, Figure 2f). However, for CoCl_2_ and Simvastatin, the result of the phenotypic adaptation was different between these two chemicals, as CoCl_2_ favored phenotypes with high fatty acid storage (consistent with a hypoxic response^47,48^), while simvastatin endorsed higher mitochondrial membrane potential and lysosomal acidity (Figure 2d and 2e), consistently to what reported elsewhere for other cell types^49,50^. The other stressors tested, however, had a radically different effect on the MCF10A cell culture, effectively driving the change in organelle traits towards a specific, stress-dependent phenotype. In this, ESPRESSO provided the unique advantage of not only detecting the exposure to various stressors, but also elucidating the mechanism of action on the cell population and defining a unique fingerprint for different stressors in living cells.

**Figure 2.**
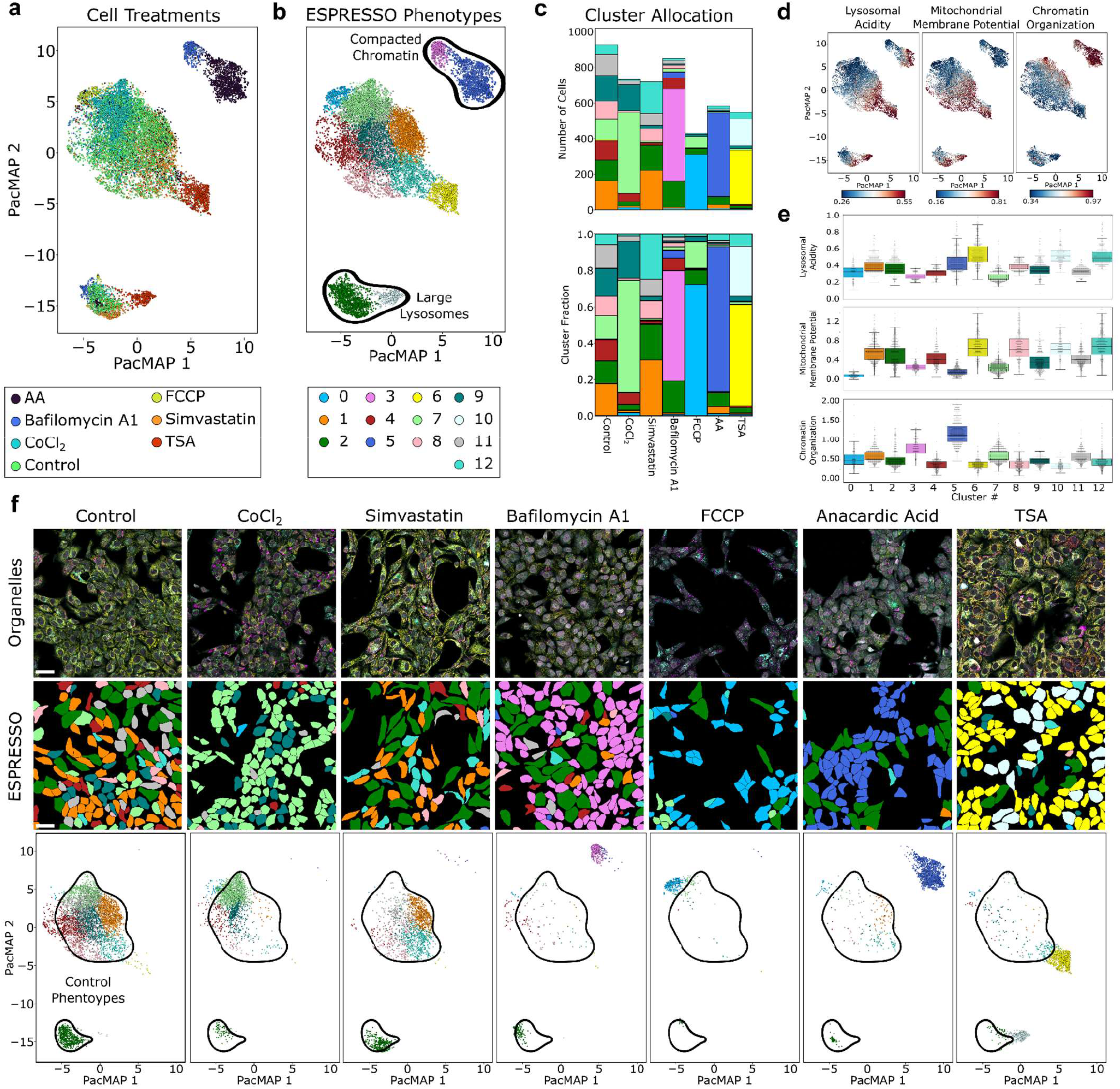
Differential ESPRESSO phenotypic response to stressor exposure. ESPRESSO PacMAP color-coded by cell treatment (a) and ESPRESSO phenotypes (b), regions highlighted in b correspond to phenotypes characterized by compacted chromatin (top) and large cells (bottom), respectively. Stacked bar plots (c) highlighting the number of cells (top) and fraction (bottom) of ESPRESSO phenotypes arranged by cell treatment. ESPRESSO PacMAPs color-coded by representative organelle properties (d) and corresponding swarm plots and box-and-whiskers plots (e). The black central line in the box-and-whiskers plot denotes the median value, the box extends between the upper and lower quartile and the whiskers extend between the minimum and maximum value, excluding outliers. Organelle properties are calculated as the center of mass of the intensity distribution, cross- or auto-correlation curves, as described in the Methods section. Representative regions of interest (f) showing the unmixed organelles images (top, white: DNA, yellow: mitochondria, cyan: lysosomes, magenta: lipid droplets), the cell masks color-coded by ESPRESSO phenotype (middle) and corresponding ESPRESSO PacMAPs (bottom, regions highlight phenotypes present in control conditions) scalebars in f are 50 μm. AA: Anacardic Acid, FCCP: Carbonyl cyanide-p-trifluoromethoxyphenylhydrazone, TSA: Trichostatin A, LD: Lipid Droplet, Mito: Mitochondria

### ESPRESSO time-resolved phenotypic evolution during keratinocyte differentiation

Moving forward from stationary analysis of cell states, we turned to two major cellular processes that are intrinsically time-resolved: cell differentiation and macrophage polarization.

Cell differentiation characterizes development in its evolution from single cells to complex living organisms, and is a dynamic process that involves reprogramming of gene expression profiles. Such reprogramming defines the boundaries of a cell’s function and it’s a highly regulated process that has been shown to have repercussions on the organelle landscape on various cell types. In skin, keratinocytes undergo a continuous process of differentiation that begins in the basal layer, the deepest of the layers in the epidermis. As they divide, newly formed keratinocytes undergo progressive differentiation as they approach the cornified layer, the outermost layer, where they ultimately reach their terminal differentiation state, culminating in regulated cell death^51^. *In vitro*, keratinocytes can be grown in culture in low-calcium medium and their differentiation can be triggered by supplementing calcium. As described in Figure 3a, we seeded keratinocytes in either control conditions or supplemented with 1.2 mM of Calcium to induce differentiation. We imaged them over the course of 22.5 hours at 30-minute intervals, collecting information on 73,430 total cells. ESPRESSO reports striking phenotypic differences between keratinocytes in control (Figure 3b) and calcium-switch (Figure 3c) conditions, showing the emergence of four differentiation-specific ESPRESSO phenotypes, characterized by drastic morphological and functional changes in the organelle landscape (Figure 3f), such as an increase in nuclear size and lysosomal acidity in specific calcium-induced differentiation cell states, consistently with trends seen during keratinocyte differentiation *in vivo*^52,53^. Moreover, we were also able to assess the phenotype’s temporal evolution, appreciating the difference between phenotypes that arise quickly and others that manifest at a later stage (Figure 3d and 3e), suggesting heterogeneity in the differentiation response. Interestingly, the phenotypic temporal evolution is also accompanied by a cellular spatial rearrangement. In fact, the cell culture is no longer composed of isolated cells, and keratinocytes tend to organize in patches (Figure 3g), suggesting that cell-cell contacts are increased upon calcium-induced differentiation and mimicking *in vivo* trends^54^. As phenotypic changes during keratinocyte differentiation span across both space and time, this application perfectly showcases the power of ESPRESSO spatiotemporal omics in identifying not only the presence of distinct phenotypes, but also providing insights about their spatiotemporal evolution.

**Figure 3.**
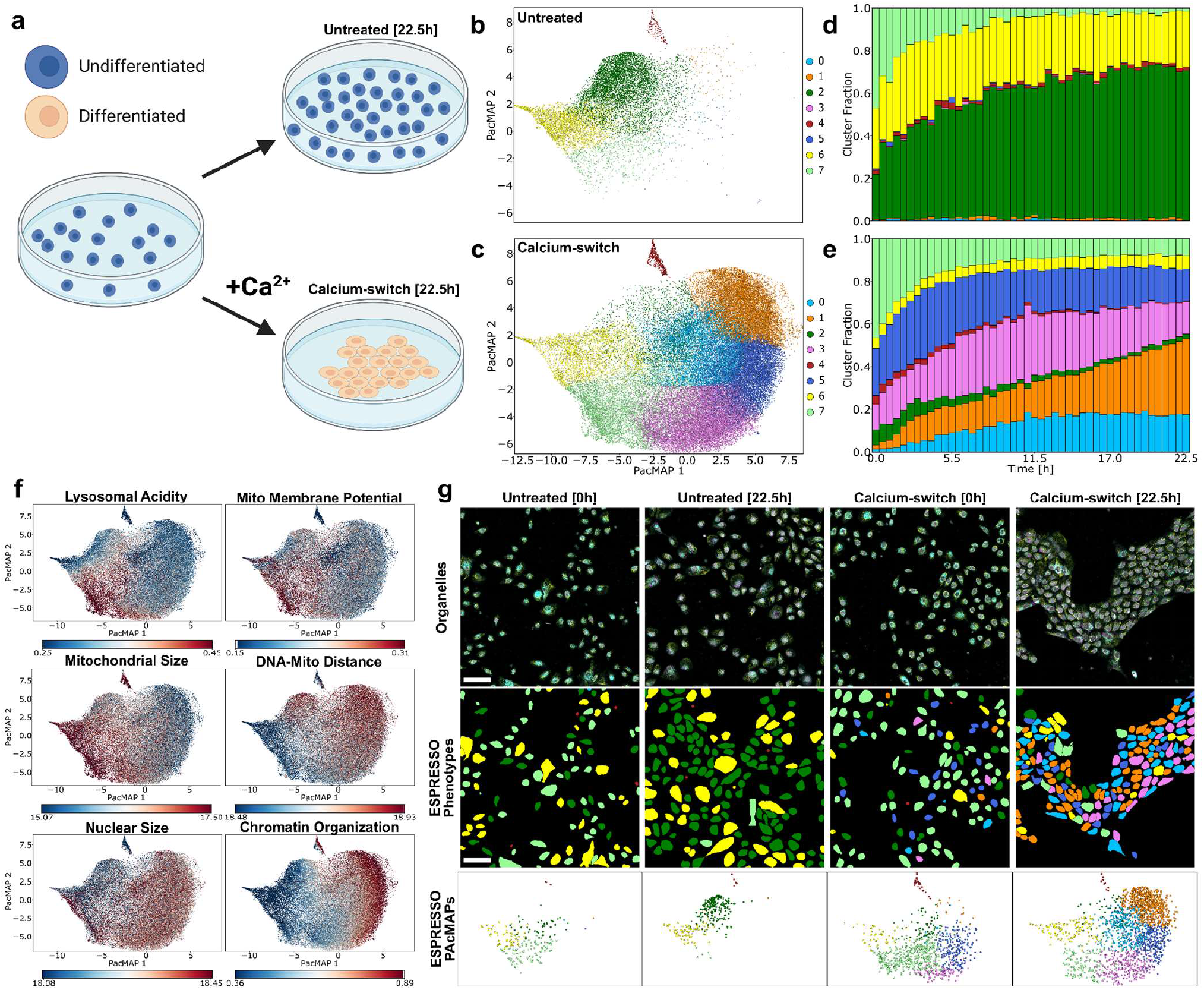
ESPRESSO phenotype evolution during keratinocyte differentiation. Schematic representation of experimental workflow (a), N/TERT-2G keratinocytes are plated and imaged for 22.5 hours in growth conditions (top) or in medium supplemented with 1.2 mM Ca2+ to induce differentiation via calcium switch. ESPRESSO PacMAPs of 73,430 total cells in untreated (b) and calcium-switch (c) conditions, color-coded according to the ESPRESSO phenotypes. Stacked bar plot showing the cluster fraction as a function of time for keratinocytes in untreated (d) and calcium-switch (e) conditions. ESPRESSO PacMAPs color-coded by representative organelle properties (f), calculated as the center of mass of the intensity distribution, cross- or auto-correlation curves, as described in the Methods section. Representative regions of interest (g) showing the unmixed organelles images (top, white: DNA, yellow: mitochondria, cyan: lysosomes, magenta: lipid droplets), the cell masks color-coded by ESPRESSO phenotype (middle) and corresponding ESPRESSO PacMAPs (bottom) scalebars in f are 100 μm. Mito: Mitochondria

### ESPRESSO phenotypic evolution during macrophage M1 activation correlates with gene expression

Next, we sought to understand if ESPRESSO phenotypes can be correlated with gene expression. To answer this question, we focused on phenotyping the process of macrophage polarization. Macrophages^55^ are immune cells targeting pathogens as well as bacteria or debris and are characterized by a process named polarization, trigger by and inflammatory response, and leads to the acquisition of two broad phenotypes defining the categories of pro-inflammatory (or M1-polarized) and anti-inflammatory (or M2-polarized) macrophages^56^. M1 polarization can be achieved *in vitro* by exposing the cells to lipopolysaccharide (LPS), a component of the outer membrane of gram-negative bacteria. LPS is recognized as an inflammatory agent by macrophages, initiating M1 polarization as a result. M1 polarization is characterized by overexpression of numerous cytokines and chemokines, together with enzymes and other proteins that prepare the cell to counteract a potential infection. Among them, iNOS (inducible nitric oxide synthase) plays a crucial role by producing nitric oxide (NO) and its expression is regulated by the *NOS2* gene^57^. Here, we utilized an immortalized mouse macrophage cell line (RAW264.7-NOS2^39^) stably transduced with a gene expression reporter for the *NOS2* gene. Upon *NOS2* gene expression, the cell line also expresses CeNL, a bioluminescent reporter comprising a fluorescent protein (mTurquoise2): the intensity of the fluorescent reporter correlates directly to the *NOS2* expression. For this experiment, we seeded transduced RAW264.7-NOS2 macrophages in growth conditions and in presence of LPS (Figure 4a) and imaged them for 24 hours in 30-minute intervals. Importantly, we simultaneously acquired the ESPRESSO phenotype as well as mTurquoise2 fluorescence intensity, allowing for the concurrent detection of organelle properties and *NOS2* gene expression. Data analysis identifies three clusters (0, 6 and 9) as ESPRESSO phenotypes that are exclusive or enriched after 24 hours of LPS exposure. These clusters are characterized by chromatin reorganization and an increase in lysosomal acidity and mitochondrial membrane potential, effects also reported in the literature. Importantly, we quantify the gene reporter fluorescence intensity levels and found all three phenotypes to significantly upregulate *NOS2* expression (Figure 4 d and g), providing a direct link between gene expression and the organelle reorganization.

**Figure 4.**
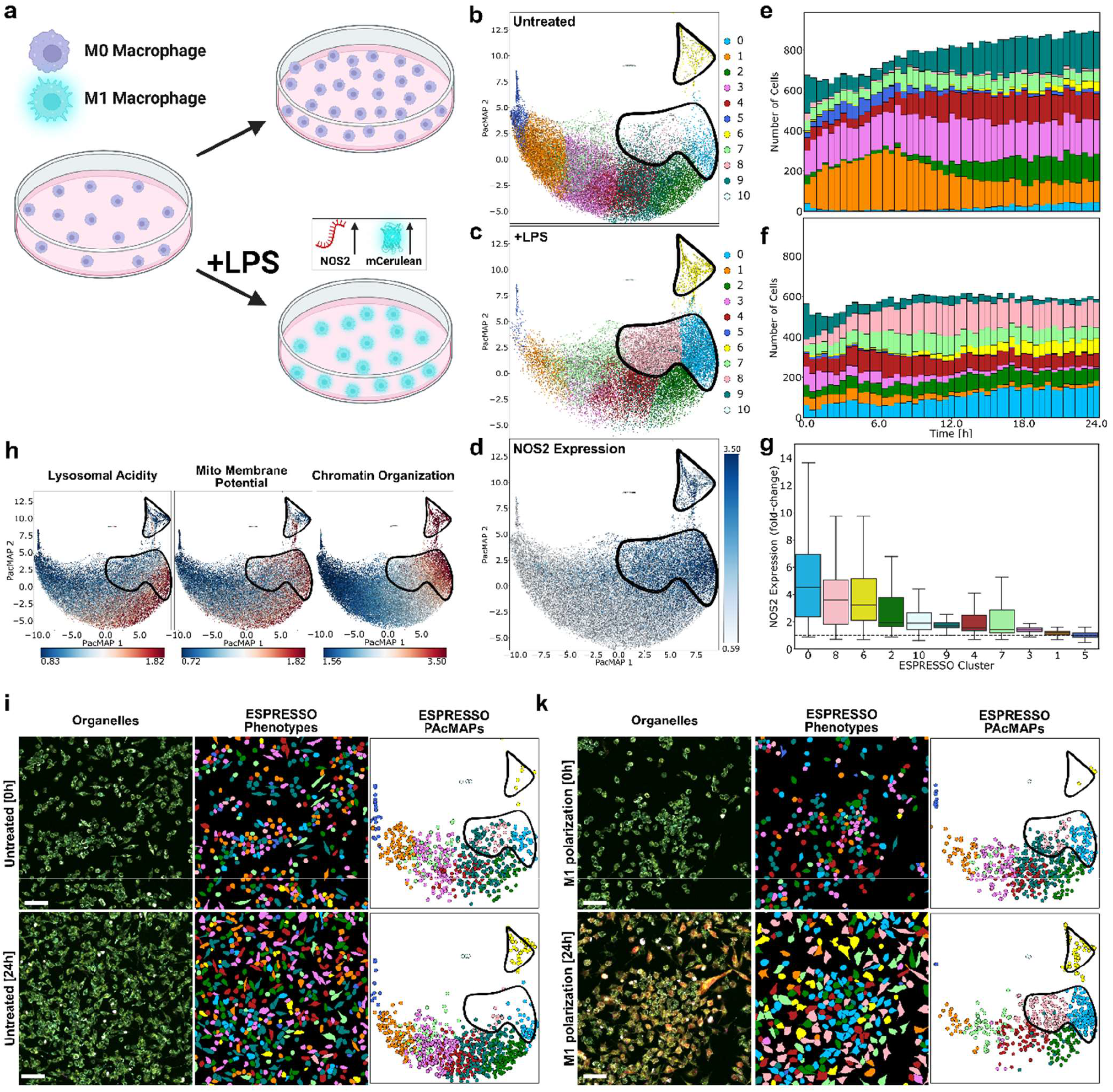
ESPRESSO phenotypic evolution during macrophages M1 polarization correlates with gene expression. Schematic representation of experimental workflow (a), RAW 264.7 macrophages transduced with a NOS2-mCerulean gene expression reporter are plated and imaged for 24 hours in growth conditions (top) or in medium supplemented with 500 ng/ml LPS to induce M1 polarization. ESPRESSO PacMAPs of 67,381 total cells in untreated (b) and M1 polarization (c) conditions, color-coded according to the ESPRESSO phenotypes. ESPRESSO PacMAP (d) color-coded according to NOS2 gene expression, calculated as the center of mass of the mCerulean intensity distribution. Stacked bar plot showing the number of cells per cluster as a function of time for macrophages in untreated (e) and M1 polarization (f) conditions. Box-and-whiskers plot (g) showing the distribution of NOS2 expression per cluster as fold-change with respect to the cluster with the least expression. Black dotted line passing by 1 serves as visual guide. The central black line identifies the median while the extension of the box identifies the upper and lower quartiles, the whiskers extend to the minimum and maximum value, excluding outliers. ESPRESSO PacMAPs color-coded by representative organelle properties (h), calculated as the center of mass of the intensity distribution as described in the Methods section. Representative regions of interest (i, k) at 0 (top) and 24 hours (bottom)showing the unmixed organelles images (left, white: DNA, yellow: mitochondria, cyan: lysosomes, magenta: lipid droplets, red: mCerulean), the cell masks color-coded by ESPRESSO phenotype (center) and corresponding ESPRESSO PacMAPs (right) scalebars in f are 50 μm. Mito: Mitochondria

### Phenotypic response in 3D cultures: Tumor spheroids invasion and response to stressors

So far, we have applied ESPRESSO phenotyping to static or longitudinal phenotyping of cell types, stressor exposure, differentiation and polarization in 2D cell cultures. Many biological systems, though, are not well recapitulated *in vitro* in 2D cultures, but rather in complex 3D systems^58–60^. An example of these systems is triple-negative breast cancer (TNBC), a highly aggressive class of breast cancer and resistant to chemotherapy due to the loss of three major chemotherapy targets (estrogen receptor, progesterone receptor and HER2^61^c). In breast cancer studies, tumor spheroids and organoids have been shown to better recapitulate the cell types found *in vivo*, making them an invaluable tool in the battle against cancer^62,63^. We sought to demonstrate that ESPRESSO can be applied longitudinally to complex 3D cultures and can be used to assess phenotype-specific responses to stressors in 3D tumor spheroids. To this aim, we generated tumor spheroids from triple-negative breast cancer cells (MDA-MB-231) and subsequently embedded them in a collagen matrix, similarly to our previous work^22,64^. Over time, cells at the surface of the spheroids undergo a metabolic switch and begin invading the collagen matrix, an event occurring at approximately 12 to 16 hours post-embedding. In order to capture the phenotypic evolution of single cells in TNBC spheroids, we performed a 16-hour volumetric time-lapse from 9 to 25 hours post-embedding. The arising of two new clusters (0 and 3) can be appreciated in Figure 5 d and e. We postulate that these phenotypes are the collagen-invading phenotypes, as also supported by the fact that cells in these clusters localize farther from the spheroid center (Figure 5 c). This analysis can precisely identify the characteristics of collagen-invading phenotypes, providing tools to specifically target them.

**Figure 5.**
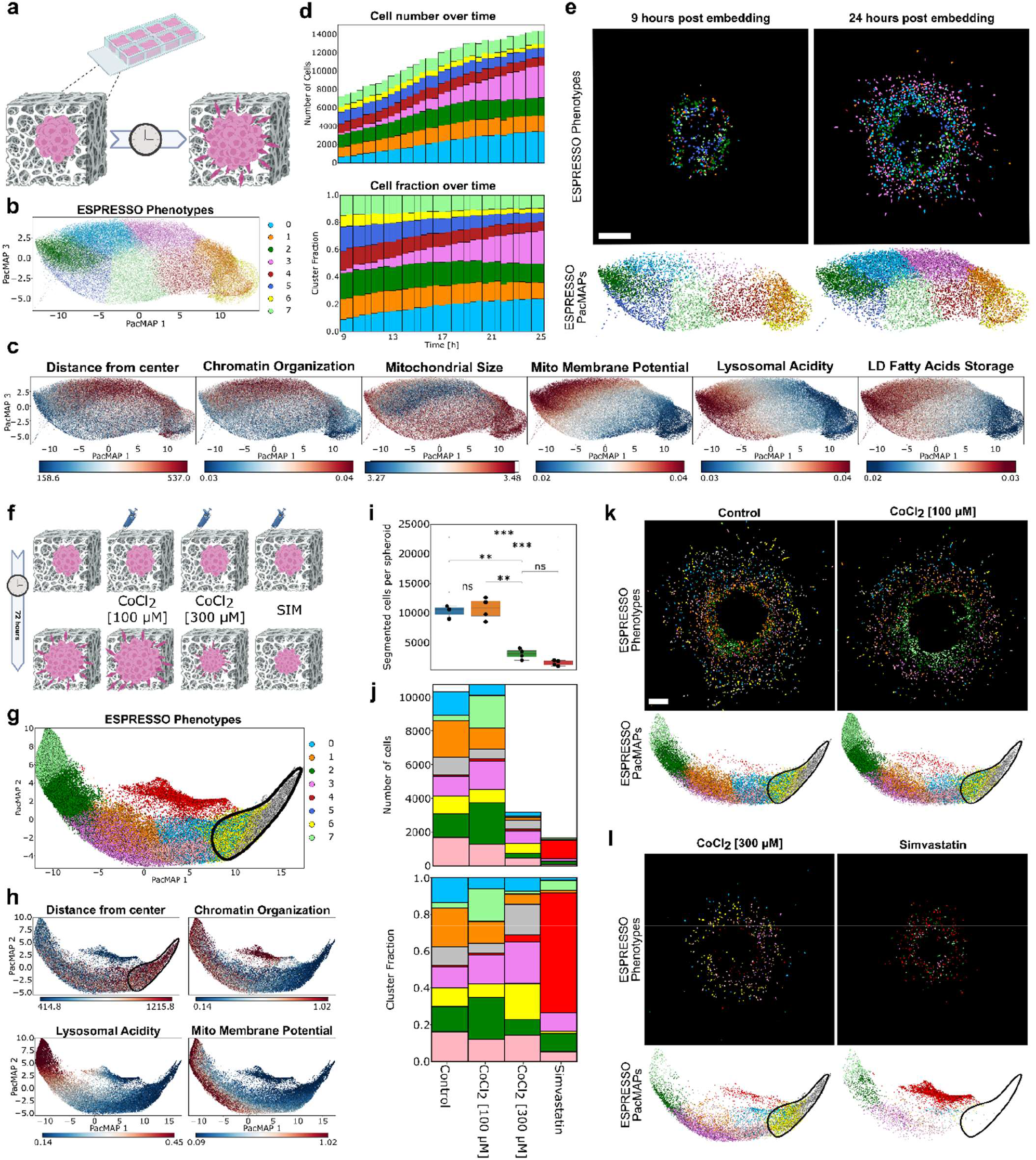
ESPRESSO in 3D tumor spheroids: phenotypic evolution and response to stressors. Schematic representation of experimental workflow for the phenotypic evolution experiment (a), MDA-MB-231 spheroids are generated, embedded in collagen and imaged for 16 hours (9-25 hours post embedding) in growth conditions. ESPRESSO PacMAP (b) of 375,964 total cells obtained from 3 spheroids and color-coded according to the ESPRESSO phenotypes. ESPRESSO PacMAPs color-coded by representative organelle properties (c.), calculated as the center of mass of the intensity distribution or auto-correlation curves, as described in the Methods section. Stacked bar plot showing the number of cells (d, top) and cluster fraction (d, bottom) as a function of time. Representative images and PacMAPs (e) showing the cell masks color-coded by ESPRESSO phenotype (top) and corresponding ESPRESSO PacMAPs (bottom) for a tumor spheroid at 9 (left) and 25 hours (right) post embedding. scalebars in e are 250 μm. Schematic representation of experimental workflow for the stressor experiment (f), MDA-MB-231 spheroids are generated, embedded in collagen and imaged after 72 hours in growth conditions (control) or in culturing medium containing 100 μM CoCl2, 300 μM CoCl2 or Simvastatin. ESPRESSO PacMAP (g) of 97,150 total cells obtained from 4 spheroids per condition and color-coded according to the ESPRESSO phenotypes. ESPRESSO PacMAPs color-coded by representative organelle properties (h), calculated as the center of mass of the intensity distribution as described in the Methods section. Number of segmented cells per spheroid (i) for the different conditions, statistical significance calculated as Student’s t-test with Bonferroni correction. (ns: p-value>0.05, **: 0.001<p-value<0.01, ***: 0.0001<p-value<0.001). Stacked bar plot showing the number of cells (j, top) and cluster fraction (j, bottom). Representative images and PacMAPs (k and l) showing the cell masks color-coded by ESPRESSO phenotype (top) and corresponding ESPRESSO PacMAPs (bottom) for representative tumor spheroids in control (k, left), CoCl2 [100 μM] (k, right), CoCl2 [200 μm] (l, left) and Simvastatin (l, right). scalebars in e are 250 μm. Mito: Mitochondria, LD: Lipid Droplets

A common parameter used to assess the viability of a spheroid culture is to evaluate the size (or number of cells) as a function of time or concentration of a specific treatment^65^. Nonetheless, for a more comprehensive evaluation of treatment response, it is essential to consider the nature of the targeted cells. Therefore, we emphasize the critical importance of analyzing the phenotypic composition of the spheroid. Indeed, we will show that ESPRESSO can identify whether or not the collagen-invading phenotypes in 3D TNBC organoids are targeted by an external treatment.

To achieve this result, we exposed the spheroids to CoCl_2_ and Simvastatin for 72 h and mapped their ESPRESSO phenotype. We used two different concentrations of CoCl_2_ with little (100 μM) or high (300 μM), reported to have different effects on spheroid growth^66^, as reflected by the cell number per spheroid (Figure 5i) being unaffected for 100 μM and dramatically decreasing for 200 μM CoCl_2_. Additionally, we considered simvastatin at a concentration causing a similar reduction in the spheroid cell number^67^. First, we identified the collagen-invading phenotypes (2, 5 and 8) as the phenotypes that characterize cells that have migrated farther from the center of the spheroid. Then, we can dig deeper and notice that CoCl_2_ has an effect on the spheroids, while not affecting the number of cells per spheroid. This effect is mostly localized to cells closer to the surface of the spheroid, where cells undergo a major phenotypic change towards highly acidic lysosomes, while the collagen-invading phenotypes are not affected. Indeed, even at a cytotoxic concentration of 200 μM, we can notice a major phenotypic change on the surface of the spheroid, while the collagen-invading phenotypes are still present in a high percentage, suggesting a resistance of collagen-invading phenotypes to hypoxic conditions, as reported also elsewhere. On the other hand, while also reducing the number of cells per spheroid, simvastatin exposure results in a massive population shift towards a unique phenotype (cluster 4) and, more importantly, in a drastic reduction of the collagen-invading phenotypes. While the beneficial effect of simvastatin in the treatment of breast cancer has been investigated in the past and recently, with ESPRESSO we can show that the mechanism underlying this outcome relies on the ability of simvastatin to target collagen-invading TNBC phenotypes *in vitro*, introducing a unique phenotype that is underrepresented in both control and hypoxic conditions

## Discussion

While current omics techniques offer valuable snapshots of cellular states, they often fail to capture the dynamic nature of biological processes. This is particularly true for metabolic transitions, where cells undergo rapid reprogramming of gene expression, protein activity, and metabolite usage. To fully understand these transitions and their role in health and disease, time-resolved omics data with high temporal resolution is crucial. While methods that allow time-resolved phenotyping exist, they are often limited to particular processes^68^, require heavy training^69^, require genetic manipulation^68^ and/or are not applicable to 3D models^69,70^. Indeed, a high-dimensional phenotyping technique that is applicable at the single-cell level, time-resolved and that doesn’t require genetic manipulation would be game-changing in omics technology, with important repercussions in biology and medicine. Such data would allow researchers to track the step-by-step changes occurring during the transition, revealing key regulatory points and potential therapeutic targets. The development of faster, more sensitive omics techniques is therefore essential for unlocking the secrets of these dynamic metabolic shifts. However, despite the organelle landscape’s importance, consistent, broadly applicable methods for reliably extracting morphological and functional information from organelles, especially in a time-resolved manner, are lacking. This stems from several technical and technological barriers: the lack of stable, organelle-specific fluorescent dyes suitable for live-cell imaging hinders longitudinal studies; the limited number of snapshot multicolor technologies results in long integration times and high phototoxicity; and existing profiling methods are typically use-specific and difficult to generalize to other biological samples.

This work introduces ESPRESSO (Environmental Sensor Phenotyping RElayed by Subcellular Structures and Organelles), a spatiotemporally resolved, high-dimensional, single-cell technique aimed at improving our understanding of the organelle landscape in health and disease. As omics technologies have progressed from bulk, to single-cell, to spatial analyses, the need to access dynamic interactions within cellular systems has become increasingly important. ESPRESSO fills this critical gap by focusing on organelles and subcellular compartments, essential cellular components that adapt their morphological and functional properties upon changes in cellular homeostasis and response to stimuli. Traditionally, organelle properties have been linked to specific cell states, though no methods that can comprehensively report on such properties across cellular systems is available to researchers. ESPRESSO overcomes numerous obstacles by providing an innovative framework that combines stable, organelle-specific fluorescent dyes with advanced imaging techniques and sophisticated data analysis tools, ultimately enabling continuous assessment of a cell’s state for over 24 hours, at the single-cell level and resolved in both space and time. The significance of ESPRESSO is demonstrated through a series of experimental settings across various cellular systems. In numerous key biological phenotyping tasks, ranging from discerning cell types to the study of phenotypic adaptation to stressors, keratinocyte differentiation and macrophage polarization, ESPRESSO unveils intricate phenotypic changes and shows correlation with gene expression. Moreover, its applicability extends to complex 3D cultures, such as tumor spheroids, where it elucidates phenotype-specific responses to environmental conditions as well as drug exposure that elude current phenotyping methods. Strikingly, ESPRESSO provides not only a snapshot of cellular states but also a dynamic understanding of their temporal evolution. By capturing spatiotemporal changes in organelle landscapes, ESPRESSO offers unprecedented insights into cellular plasticity, paving the way for new discoveries in biology and potential therapeutic interventions in various disease states. In conclusion, ESPRESSO represents a paradigm shift in phenotyping techniques, providing the research community with a novel, powerful tool to explore the intricate workings of complex cellular systems with unrivaled detail and precision.

## Methods

### Cell culture

RAW264.7-NOS2 cells and MDA-MB-231 cells were cultured in DMEM medium (DMEM, high glucose, with L-glutamine, GenClone), supplemented with 10% v/v FBS (heat inactivated FBS, GenClone) and 1% v/v penicillin/streptomycin solution 100× (10,000 units of penicillin and 10 mg ml−1 streptomycin in 0.85% saline solution, GenClone). RAW264.7-NOS2 cell line is stably expressing the Nano-lantern CeNL under the NOS2 promoter as described in our previous work^39^, which consist of a luminophore (Nluc) linked to a fluorescent protein (mTurquoise2)^71^. MCF10A cells were cultured in DMEM/F12 medium (Gibco), supplemented with 5% v/v Horse Serum (heat inactivated, Invitrogen), Epidermal Growth Factor (Peprotech), Hydrocortisone (Sigma), Cholera Toxin (Sigma), Insulin from bovine pancreas (Sigma), 1% v/v penicillin/streptomycin solution 100× (10,000 units of penicillin and 10 mg ml−1 streptomycin in 0.85% saline solution, GenClone). N/TERT-2G keratinocytes^38^ (a gift from Jos P. H. Smits, Radboud University Medical Center, Nijmegen, The Netherlands) were cultured in Keratinocyte SFM (1X) medium (Gibco), supplemented with 50 μg/mL Bovine Pituitary Extract (BPE) (Gibco) and 5 ng/mL Human Recombinant EGF (Gibco) following manufacturer instructions, and 1% v/v penicillin/streptomycin solution 100× (10,000 units of penicillin and 10 mg ml−1 streptomycin in 0.85% saline solution, GenClone). All cell lines were cultured in a 37 °C and 5% CO2 incubator.

### Labeling

LysoTracker™ Green DND-26 (Lysotracker Green, ThermoFisher Scientific L7526), Tetramethylrhodamine methyl ester (TMRM, Cayman Chemical 21437), SMCy5.5 (Bio-Techne #7295) and SiR-DNA (SiR-Hoechst, Cytoskeleton Inc.CY-SC007) were used throughout this work. A 1000x mixture of these dyes (Lysotracker Green 200 μM, TMRM 100 μM, SMCy5.5 3 μM and SiR-Hoechst 670 μM) was prepared in DMSO and stored for a maximum of 4 weeks. The mixture was applied at 1x to cells upon plating, diluted with the culturing medium, and at the moment of administering the treatments.

### 2D sample preparation

Cells were seeded in 8-well chambers (Cellvis) in their culture medium containing 1X dye mixture. After 24 hours half of the medium was replaced with fresh medium containing 1X dye mixture. MDA-MB-231, MCF10A and N/TERT-2G cells were seeded at 30,000 cells/well, RAW264.7-NOS2 cells were seed at 50,000 cells/well.

For the stressor-exposure experiments, MCF10A cells were seeded in 8-well imaging chambers (Cellvis) at 30,000 cells/well in culture medium containing 1X dyes mixture. The day after, the medium was replaced with medium containing 1X dyes mixture for control and medium containing 1X dyes mixture + treatments, with the following final concentrations: Anacardic Acid (Sigma-Aldrich) 60 μM, Trichostatin A (Sigma-Aldrich) 1 μM, Bafilomycin A1 (Cayman Chemical) 50 nM, Carbonyl cyanide 4- (trifluoromethoxy)phenylhydrazone - FCCP (Sigma-Aldrich) 1.25 μM, Cobalt chloride CoCl_2_ 400 μM (Sigma-Aldrich), Simvastatin (Sigma-Aldrich) 20 μM. Cells were incubated for 24 hours with the treatments before imaging. Upon thawing, MCF10A cells were split in three flasks and passed in parallel for at least two passages, so that each flask constitutes a biological replicate. Each condition described was tested on each of the replicates, with the exclusion of FCCP and Bafilomycin A1 that was tested on a single replicate.

For macrophages polarization experiments, RAW264.7-NOS2 cells were seed at 50,000 cells/well in culture medium containing 1X dyes mixture. The day after, half of the medium was replaced with medium containing 1X dyes mixture for control and medium containing 1X dyes mixture and Lipopolysaccharide (LPS) Solution (Invitrogen) with a final concentration of 500 ng/mL for the polarization treatment.

For the keratinocytes calcium switch experiments, N/TERT-2G cells were seeded in 8-well imaging chambers (Cellvis) at 30,000 cells/well in culture medium containing 1X dyes mixture. The day after, half of the medium was replaced with medium containing 1X dyes mixture for control and medium containing 1X dyes mixture and Calcium Chloride (Gibco) at a final concentration of 1.2 mM for the calcium-switch treatment. Upon thawing, N/TERT-2G cells were split in three flasks and passed in parallel for at least two passages, so that each flask constitutes a biological replicate. The control dataset was acquired on a single biological replicate in three technical replicates (multiple position within the same well) while the calcium-induced differentiation was performed on all three biological replicates in three technical replicates (multiple position within the same well).

### 3D sample preparation

MDA-MB-231 spheroids were formed starting from a culture of 10,000 cells per spheroid mixed with their medium and a 2% v/v Cultrex UltiMatrix Reduced Growth Factor Basement Membrane Extract (BioTechne, R&D systems) solution in 96-well round-bottom plates (Corning Costar Ultra-Low Attachment Multiple Well Plate), similarly to our previous work^22,64^. After a brief centrifugation (5 min at 1,000 rpm.) and one day of incubation (37 °C, 5% CO2), the spheroids labeled with the dye mixture. The following day the spheroids were transferred in eight-well chambers (Cellvis) and embedded in a 2.0 mg ml−1 collagen Type-1 gel matrix (Corning Collagen I, High concentration, Rat Tail, 100 mg). Medium containing the dyes mixtures was added on top of the matrix after collagen polymerization and incubated at 37 °C and 5% CO2 until the moment of imaging.

For the time lapse experiments, four different spheroids were prepared and imaged. For the data collected 72 hours after embedding, 4 different spheroids per condition were prepared and imaged. At the moment of medium and dyes addition on top of the polymerized collagen matrix, the following compounds were added: Simvastatin (Sigma-Aldrich) final concentration 20 μM, Cobalt chloride (CoCl_2_, Sigma-Aldrich) final concentration 100 μM or 300 μM.

### Hyperspectral Imaging and unmixing

Imaging was performed with a Zeiss LSM880 with Spectral (Quasar) detector. ESPRESSO images were acquired with a 63x/1.4 NA Oil objective (Zeiss) with a variable number of tiles, each tile consisting of a 512×512 image with 120 nm pixel size, and spheroid measurements were acquired with a 40x/0.75 NA Air objective (Zeiss). Two dichroic mirrors (‘MBS 488/561/633’ and ‘MBS -405’) were selected to allow excitation with 488, 561, 633 and/or 405 nm laser lines. All measurements in 2D were taken with both the 488 nm (0.1%, corresponding to 0.2 μW) and the 633 nm (1%, corresponding to 4 μW) laser lines, macrophage polarization measurements also include excitation with 405 nm laser (0.2 %, corresponding to 5.5 μW) in order to excite the gene expression reporter. All measurements in spheroids were taken with both the 488 nm (0.5%, corresponding to 6.9 μW) and the 633 nm (1.5%, corresponding to 7.7 μW) laser lines. Laser power was measured after the objective using a power meter set for each specific wavelength (Thorlabs). Images are saved as Zeiss .lsm files. For spectral unmixing, average emission spectra from MDA-MB-231 cells labeled with the individual dyes have been imaged and saved in a database. During processing, the spectra are loaded and transformed in spectral coordinates (up to the 2nd harmonic, allowing up to 5-color unmixing) and used as pure components for linear unmixing. Negative values in the unmixed images are set to 0. The raw (.lsm) images after unmixing are saved as .npy files.

### CARE denoising training

CARE network has been trained on a total of 128 noisy and 128 high SNR 512×512 images of MDA-MB-231 and MCF10A cells. Images were acquired sequentially, with the noisy image acquired first and the second acquired with the same parameters but averaging 16 times each line for higher SNR. For each image, the mean intensity of the high SNR image was scaled to the mean intensity of the corresponding low SNR image, and Poisson noise is artificially added to degrade it. The degraded high SNR image is used for training as the organelles are in the exact same position as in the non-degraded high SNR image, that is used as ground truth. Each image is spectrally unmixed by phasor and each unmixed channel (lysosomes, mitochondria, lipid droplets and DNA) is trained separately. Training is performed on 64x64 patches, of which only the 50% brightest were considered. Then, data is split 80/20 in training/validation, and the CARE network is trained with a kernel size of 3, batch size of 8 for 250 epoch for each channel. The CARE network and the training parameters are saved in a dictionary that is used as an input in the ESPRESSO pipeline.

**Table 1:**
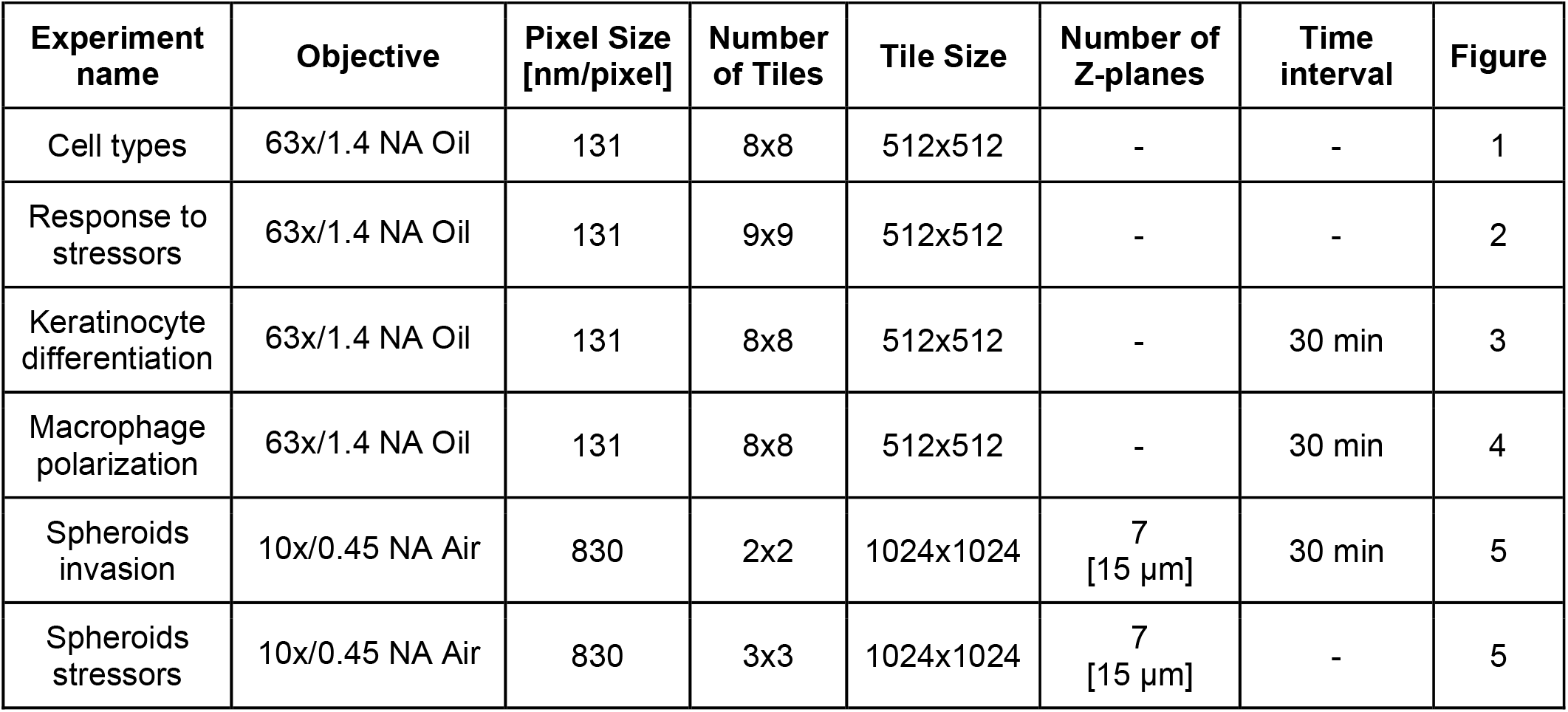
Table describing the imaging parameters for each experiment presented in this work.

### Image processing parameters

Single-cell segmentation is performed with CellPose (version 2.2.2) using the pre-trained network “cyto2”. Input images are 2-channel images consisting of the SiR-Hoechst channel (nucleus) and the sum of the other organelle channels (cytoplasm) binned 2×2, normalized to the 99th percentile and converted in uint8. Different diameters are entered for the different cell types (MDA-MB-231 (2D), N/TERT-2G and MCF10A: 175, RAW246.7-NOS2: 125, MDA-MB-231 (3D): 30).

### Feature extraction

One segmented, the organelle images for each cell are analyzed as follows. For each individual channel, the intensity distribution is calculated over 512 logarithmic bins ranging from 10^−2^ to 10^3^ and normalized so that its sum equals 1, and the ICS autocorrelation function is calculated. For each ICS autocorrelation function, the rotational average is obtained and padded with zeros to 128 pixels. For each channel combination, the ICCS image cross-correlation function is computed and analyzed in the same way as ICS. Each bin of these functions (intensity histograms, ICS and ICCS functions) are passed as features for the next analysis step, yielding a total of 3,328 features. A Pandas DataFrame of the same length of the number of segmented cells is compiled and the organelle features are stored along with the position of the cells within the image (calculated as the center of mass of the CellPose mask), the name of the file and the ID of the cell.

### Feature selection

For data analysis, multiple files are entered in the same pipeline (e.g., experimental replicates, control and treatment conditions, multiple timepoints…), are identified based on their file name and a name is assigned to each file depending on which condition it refers to. Additionally, the conditions on which to perform the feature comparison are chosen. For static experiments, all conditions are considered, while for dynamics experiments only the final time point is used for feature selection. For every feature for each pair of conditions considered for feature comparison, a Kolmogorov-Smirnov test with Bonferroni correction for multiple comparisons is performed and the p-value is stored along with the difference between the median value across all cells for each feature. A scatterplot (analog to the volcano plot for scRNA-seq) of -log_10_(p-value) as a function of the difference of the medians is drawn for functional (intensity histograms) and morphological (ICS and ICCS functions) features and the user define a range of p-value and median differences to use for dimensionality reduction. A subset of the organelle profile including only the selected features is used for dimensionality reduction.

### PacMAP and clustering

PacMAP dimensionality reduction is performed by the Python library pacmap (version 0.7.0) outputting 3 dimensions selecting random_state = 1. Only the conditions used for the feature selection are used to define the space and the cells associated to the remaining conditions are projected into that space. At this stage, a Gaussian Mixture Model (GMM) clustering step is performed with a varying number of clusters (from 2 to 16) and of initial states (from 0 to 100 in steps of 10), calculating the Davies-Bouldin Index (DBI) for each condition. After testing all conditions, the DBI as a function of the number of clusters is drawn and the number of clusters and initial state are user-defined based on the minima of the curves and the degree of detail required by the clustering.

### Data visualization

ESPRESSO PacMAPs are saved for all three dimensions as scatter plots where each point is color-coded according to either their experimental conditions, their associated cluster, together with bar plots depicting the absolute or normalized contribution of each condition or cluster. The center of mass for the organelle functions (intensity histograms, ICS and ICCS functions) is also calculated and used to color-code the ESPRESSO PacMAPs for a visual representation of the organelle properties. Finally, the segmentation masks are loaded and the mask corresponding to each cell is color-coded based on its associated cluster. For time-resolved experiments, the output is a GIF of the color-coded masks and the corresponding PacMAP, evolving as a function of time.

## References

1. Wang, Z., Gerstein, M. & Snyder, M. RNA-Seq: A revolutionary tool for transcriptomics. Nat. Rev. Genet. 10, (2009).

2. Cui, M., Cheng, C. & Zhang, L. High-throughput proteomics: a methodological mini-review. Lab. Invest. 102, (2022).

3. Johnson, C. H., Ivanisevic, J. & Siuzdak, G. Metabolomics: Beyond biomarkers and towards mechanisms. Nat. Rev. Mol. Cell Biol. 17, (2016).

4. Lewis, S. M. et al. Spatial omics and multiplexed imaging to explore cancer biology. Nat. Methods 18, (2021).

5. Valm, A. M. et al. Applying systems-level spectral imaging and analysis to reveal the organelle interactome. Nature (2017) doi:10.1038/nature22369.

6. Jain, A. & Zoncu, R. Organelle transporters and inter-organelle communication as drivers of metabolic regulation and cellular homeostasis. Mol. Metab. 60, (2022).

7. Kim, Y. et al. Characterizing Organelles in Live Stem Cells Using Label-Free Optical Diffraction Tomography. Mol. Cells 44, (2021).

8. Ahlqvist, K. J., Suomalainen, A. & Hämäläinen, R. H. Stem cells, mitochondria and aging. Biochim. Biophys. Acta - Bioenerg. 1847, 1380–1386 (2015).

9. Jarc, E. & Petan, T. Lipid droplets and the management of cellular stress. Yale J. Biol. Med. 92, (2019).

10. Frank, M. et al. Mitophagy is triggered by mild oxidative stress in a mitochondrial fission dependent manner. Biochim. Biophys. Acta - Mol. Cell Res. 1823, (2012).

11. Mukhopadhyay, S. et al. Serum starvation induces anti-apoptotic cIAP1 to promote mitophagy through ubiquitination. Biochem. Biophys. Res. Commun. 479, (2016).

12. Cruz, A. L. S., Barreto, E. de A., Fazolini, N. P. B., Viola, J. P. B. & Bozza, P. T. Lipid droplets: platforms with multiple functions in cancer hallmarks. Cell Death Dis. 11, (2020).

13. Rambold, A. S., Cohen, S. & Lippincott-Schwartz, J. Fatty acid trafficking in starved cells: Regulation by lipid droplet lipolysis, autophagy, and mitochondrial fusion dynamics. Dev. Cell 32, (2015).

14. Ramosaj, M. et al. Lipid droplet availability affects neural stem/progenitor cell metabolism and proliferation. Nat. Commun. 12, (2021).

15. Hershey, B. J., Vazzana, R., Joppi, D. L. & Havas, K. M. Lipid Droplets Define a Sub-Population of Breast Cancer Stem Cells. J. Clin. Med. 9, 87 (2019).

16. Loeffler, D. et al. Asymmetric organelle inheritance predicts human blood stem cell fate. Blood 139, (2022).

17. Bray, M. A. et al. Cell Painting, a high-content image-based assay for morphological profiling using multiplexed fluorescent dyes. Nat. Protoc. 11, (2016).

18. Novikova, S. et al. Nuclear Proteomics of Induced Leukemia Cell Differentiation. Cells 11, 3221 (2022).

19. Chikte, S., Panchal, N. & Warnes, G. Use of LysoTracker dyes: A flow cytometric study of autophagy. Cytometry A 85, (2014).

20. Scaduto, R. C. & Grotyohann, L. W. Measurement of mitochondrial membrane potential using fluorescent rhodamine derivatives. Biophys. J. 76, (1999).

21. Cutrale, F. et al. Hyperspectral phasor analysis enables multiplexed 5D in vivo imaging. Nat. Methods (2017) doi:10.1038/nmeth.4134.

22. Scipioni, L., Rossetta, A., Tedeschi, G. & Gratton, E. Phasor S-FLIM: a new paradigm for fast and robust spectral fluorescence lifetime imaging. Nat. Methods 18, 542–550 (2021).

23. Fereidouni, F., Bader, A. N. & Gerritsen, H. C. Spectral phasor analysis allows rapid and reliable unmixing of fluorescence microscopy spectral images. Opt. Express (2012) doi:10.1364/oe.20.012729.

24. Cutrale, F., Salih, A. & Gratton, E. Spectral phasor approach for fingerprinting of photo-activatable fluorescent proteins Dronpa, Kaede and KikGR. Methods Appl. Fluoresc. (2013) doi:10.1088/2050-6120/1/3/035001.

25. von Chamier, L. et al. Democratising deep learning for microscopy with ZeroCostDL4Mic. Nat. Commun. 12, 2276 (2021).

26. Weigert, M. et al. Content-aware image restoration: pushing the limits of fluorescence microscopy. Nat. Methods 15, 1090–1097 (2018).

27. Stringer, C., Wang, T., Michaelos, M. & Pachitariu, M. Cellpose: a generalist algorithm for cellular segmentation. Nat. Methods 18, (2021).

28. Kolin, D. L. & Wiseman, P. W. Advances in Image Correlation Spectroscopy: Measuring Number Densities, Aggregation States, and Dynamics of Fluorescently labeled Macromolecules in Cells. Cell Biochem. Biophys. 49, 141–164 (2007).

29. Scipioni, L., Gratton, E., Diaspro, A. & Lanzanò, L. Phasor Analysis of Local ICS Detects Heterogeneity in Size and Number of Intracellular Vesicles. Biophys. J. 111, (2016).

30. Costantino, S., Comeau, J. W., Kolin, D. L. & Wiseman, P. W. Accuracy and dynamic range of spatial image correlation and cross-correlation spectroscopy. Biophys J 89, 1251–1260 (2005).

31. Oneto, M. et al. Nanoscale Distribution of Nuclear Sites by Super-Resolved Image Cross-Correlation Spectroscopy. Biophys. J. (2019) doi:10.1016/j.bpj.2019.10.036.

32. Petersen, N. O., Hoddelius, P. L., Wiseman, P. W., Seger, O. & Magnusson, K. E. Quantitation of membrane receptor distributions by image correlation spectroscopy: concept and application. Biophys J 65, 1135–1146 (1993).

33. Liang, Z., Lou, J., Scipioni, L., Gratton, E. & Hinde, E. Quantifying nuclear wide chromatin compaction by phasor analysis of histone Förster resonance energy transfer (FRET) in frequency domain fluorescence lifetime imaging microscopy (FLIM) data. Data Brief (2020) doi:10.1016/j.dib.2020.105401.

34. Lou, J. et al. Phasor histone FLIM-FRET microscopy quantifies spatiotemporal rearrangement of chromatin architecture during the DNA damage response. Proc. Natl. Acad. Sci. U. S. A. (2019) doi:10.1073/pnas.1814965116.

35. Wang, Y., Huang, H., Rudin, C. & Shaposhnik, Y. Understanding how dimension reduction tools work: An empirical approach to deciphering T-SNE, UMAP, TriMap, and PaCMAP for data visualization. J. Mach. Learn. Res. 22, (2021).

36. Fraley, C. & Raftery, A. E. Model-based clustering, discriminant analysis, and density estimation. J. Am. Stat. Assoc. 97, (2002).

37. Xiao, J., Lu, J. & Li, X. Davies Bouldin Index based hierarchical initialization K-means. Intell. Data Anal. 21, (2017).

38. Dickson, M. A. et al. Human keratinocytes that express hTERT and also bypass a p16(INK4a)-enforced mechanism that limits life span become immortal yet retain normal growth and differentiation characteristics. Mol. Cell. Biol. 20, 1436–1447 (2000).

39. Tedeschi, G. et al. Monitoring macrophage polarization with gene expression reporters and bioluminescence phasor analysis. 2024.06.10.598305 Preprint at 10.1101/2024.06.10.598305 (2024).

40. Iriondo, O. et al. Distinct breast cancer stem/progenitor cell populations require either HIF1α or loss of PHD3 to expand under hypoxic conditions. Oncotarget 6, (2015).

41. Li, Q., Ma, R. & Zhang, M. CoCl2 increases the expression of hypoxic markers HIF-1α, VEGF and CXCR4 in breast cancer MCF-7 cells. Oncol. Lett. 15, (2018).

42. Bai, F. et al. Simvastatin induces breast cancer cell death through oxidative stress up-regulating miR-140-5p. Aging 11, 3198–3219 (2019).

43. Wang, R. et al. Molecular basis of V-ATPase inhibition by bafilomycin A1. Nat. Commun. 12, 1782 (2021).

44. Dispersyn, G., Nuydens, R., Connors, R., Borgers, M. & Geerts, H. Bcl-2 protects against FCCP-induced apoptosis and mitochondrial membrane potential depolarization in PC12 cells. Biochim. Biophys. Acta - Gen. Subj. 1428, (1999).

45. Sun, Y., Jiang, X., Chen, S. & Price, B. D. Inhibition of histone acetyltransferase activity by anacardic acid sensitizes tumor cells to ionizing radiation. FEBS Lett. 580, 4353–4356 (2006).

46. Tóth, K. F. et al. Trichostatin A-induced histone acetylation causes decondensation of interphase chromatin. J. Cell Sci. 117, 4277–4287 (2004).

47. Mylonis, I. et al. Hypoxia causes triglyceride accumulation by HIF-1-mediated stimulation of lipin 1 expression. J. Cell Sci. 125, 3485–3493 (2012).

48. Munir, R., Lisec, J., Swinnen, J. V. & Zaidi, N. Lipid metabolism in cancer cells under metabolic stress. Br. J. Cancer 120, 1090–1098 (2019).

49. Jones, S. P., Teshima, Y., Akao, M. & Marbán, E. Simvastatin Attenuates Oxidant-Induced Mitochondrial Dysfunction in Cardiac Myocytes. Circ. Res. 93, 697–699 (2003).

50. Zhang, Y. et al. Simvastatin improves lysosome function via enhancing lysosome biogenesis in endothelial cells. Front. Biosci. Landmark Ed. 25, 283–298 (2020).

51. Liu, B., Zhu, F., Xia, X., Park, E. & Hu, Y. A tale of terminal differentiation: IKKα, the master keratinocyte regulator. Cell Cycle 8, 527–531 (2009).

52. Monteleon, C. L. et al. Lysosomes support the degradation, signaling, and mitochondrial metabolism necessary for human epidermal differentiation. J. Invest. Dermatol. 138, 1945–1954 (2018).

53. Gdula, M. R. et al. Remodeling of three-dimensional organization of the nucleus during terminal keratinocyte differentiation in the epidermis. J. Invest. Dermatol. 133, 2191–2201 (2013).

54. Charest, J. L., Jennings, J. M., King, W. P., Kowalczyk, A. P. & García, A. J. Cadherin-Mediated Cell–Cell Contact Regulates Keratinocyte Differentiation. J. Invest. Dermatol. 129, 564–572 (2009).

55. Wynn, T. A., Chawla, A. & Pollard, J. W. Macrophage biology in development, homeostasis and disease. Nature 496, 445–455 (2013).

56. Chen, S. et al. Macrophages in immunoregulation and therapeutics. Signal Transduct. Target. Ther. 8, 1–35 (2023).

57. Xaus, J. et al. LPS induces apoptosis in macrophages mostly through the autocrine production of TNF-α. Blood 95, 3823–3831 (2000).

58. Nayak, P., Bentivoglio, V., Varani, M. & Signore, A. Three-Dimensional In Vitro Tumor Spheroid Models for Evaluation of Anticancer Therapy: Recent Updates. Cancers 15, 4846 (2023).

59. Ziperstein, M. J., Guzman, A. & Kaufman, L. J. Breast Cancer Cell Line Aggregate Morphology Does Not Predict Invasive Capacity. PLOS ONE 10, e0139523 (2015).

60. Lee, S.-Y., Koo, I.-S., Hwang, H. J. & Lee, D. W. In Vitro three-dimensional (3D) cell culture tools for spheroid and organoid models. SLAS Discov. Adv. Life Sci. R D 28, 119–137 (2023).

61. Huang, Z., Yu, P. & Tang, J. <p>Characterization of Triple-Negative Breast Cancer MDA-MB-231 Cell Spheroid Model</p>. OncoTargets Ther. 13, 5395–5405 (2020).

62. Hofmann, S., Cohen-Harazi, R., Maizels, Y. & Koman, I. Patient-derived tumor spheroid cultures as a promising tool to assist personalized therapeutic decisions in breast cancer. Transl. Cancer Res. 11, (2022).

63. Gilazieva, Z., Ponomarev, A., Rutland, C., Rizvanov, A. & Solovyeva, V. Promising applications of tumor spheroids and organoids for personalized medicine. Cancers (2020) doi:10.3390/cancers12102727.

64. Yao, Z. et al. Multiplexed bioluminescence microscopy via phasor analysis. Nat. Methods 19, 893– 898 (2022).

65. Reynolds, D. S. et al. Breast Cancer Spheroids Reveal a Differential Cancer Stem Cell Response to Chemotherapeutic Treatment. Sci. Rep. (2017) doi:10.1038/s41598-017-10863-4.

66. A. Naveena, H. & Bhatia, D. Hypoxia Modulates Cellular Endocytic Pathways and Organelles with Enhanced Cell Migration and 3D Cell Invasion**. ChemBioChem 24, e202300506 (2023).

67. Bytautaite, M. & Petrikaite, V. <p>Comparative Study of Lipophilic Statin Activity in 2D and 3D&nbsp;in vitro Models of Human Breast Cancer Cell Lines MDA-MB-231 and MCF-7</p>. OncoTargets Ther. 13, 13201–13209 (2020).

68. Held, M. et al. CellCognition: time-resolved phenotype annotation in high-throughput live cell imaging. Nat. Methods 7, 747–754 (2010).

69. Viana, M. P. et al. Integrated intracellular organization and its variations in human iPS cells. Nature 613, 345–354 (2023).

70. Wiggins, L. et al. The CellPhe toolkit for cell phenotyping using time-lapse imaging and pattern recognition. Nat. Commun. 14, 1854 (2023).

71. Suzuki, K. et al. Five colour variants of bright luminescent protein for real-time multicolour bioimaging. Nat. Commun. 7, 13718 (2016).

