## Supplementary Figures for "ESPRESSO: Spatiotemporal omics based on organelle phenotyping"

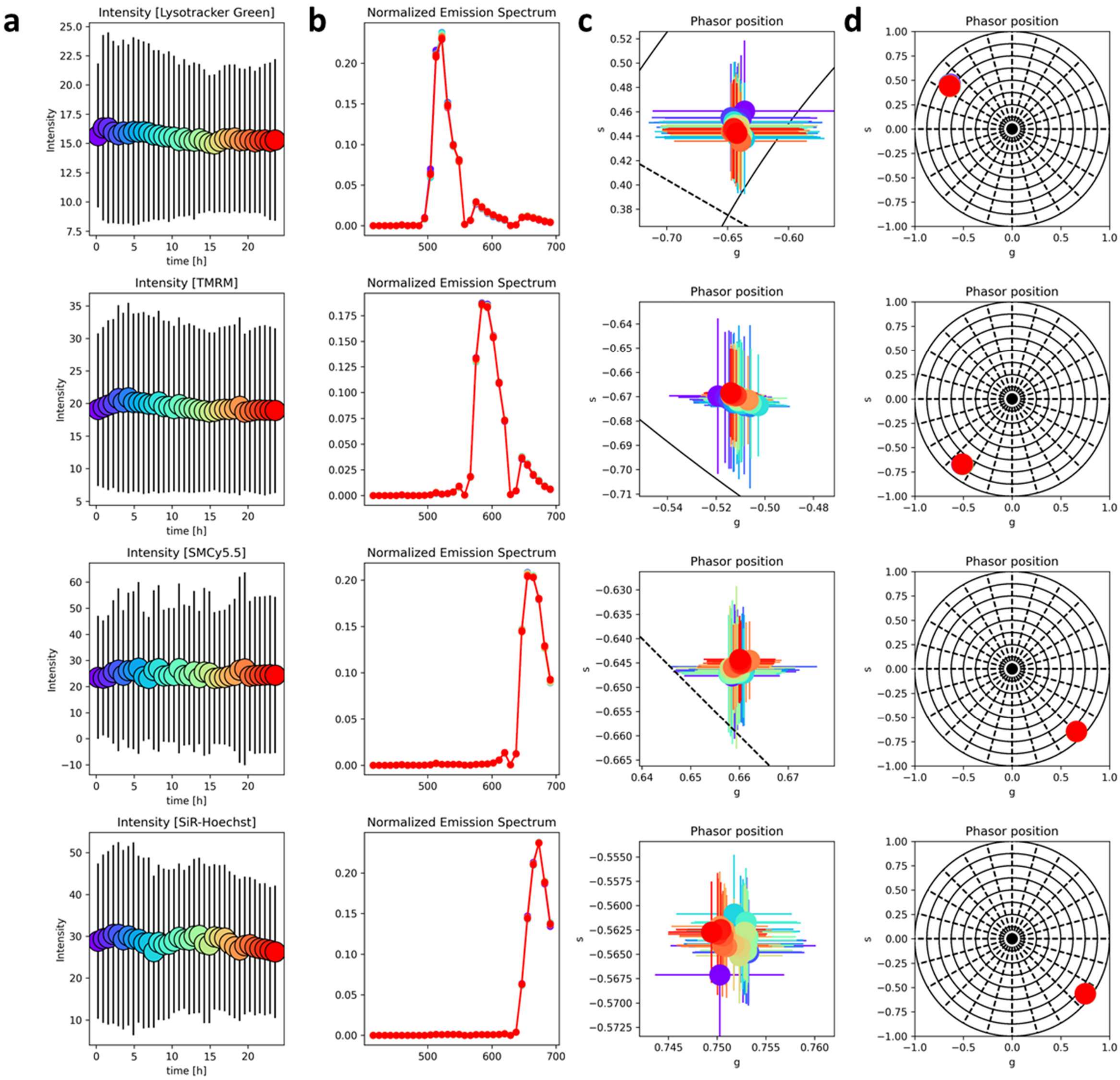


***Figure S1: Dyes stability over time***

*Average intensity (**a**), area-normalized emission spectrum (**b**), phasor position (**c**, zoomed in and **d**, full phasor plot) as a function of time for the four dyes used in this study (top to bottom): Lysotracker Green, TMRM, SMCy5.5 and SiR-Hoechst. Time is color-coded from purple (0 hours) to red (24 hours), full circles in **a** refer to the average values of pixels above threshold and error bars represent the standard deviation. Spectral in **b** are the total emission spectrum of all pixels above threshold, color-coded by time. Full circles in **c** and **d** represent the average phasor position of all pixels above threshold, and the error bars represent their standard deviation.*

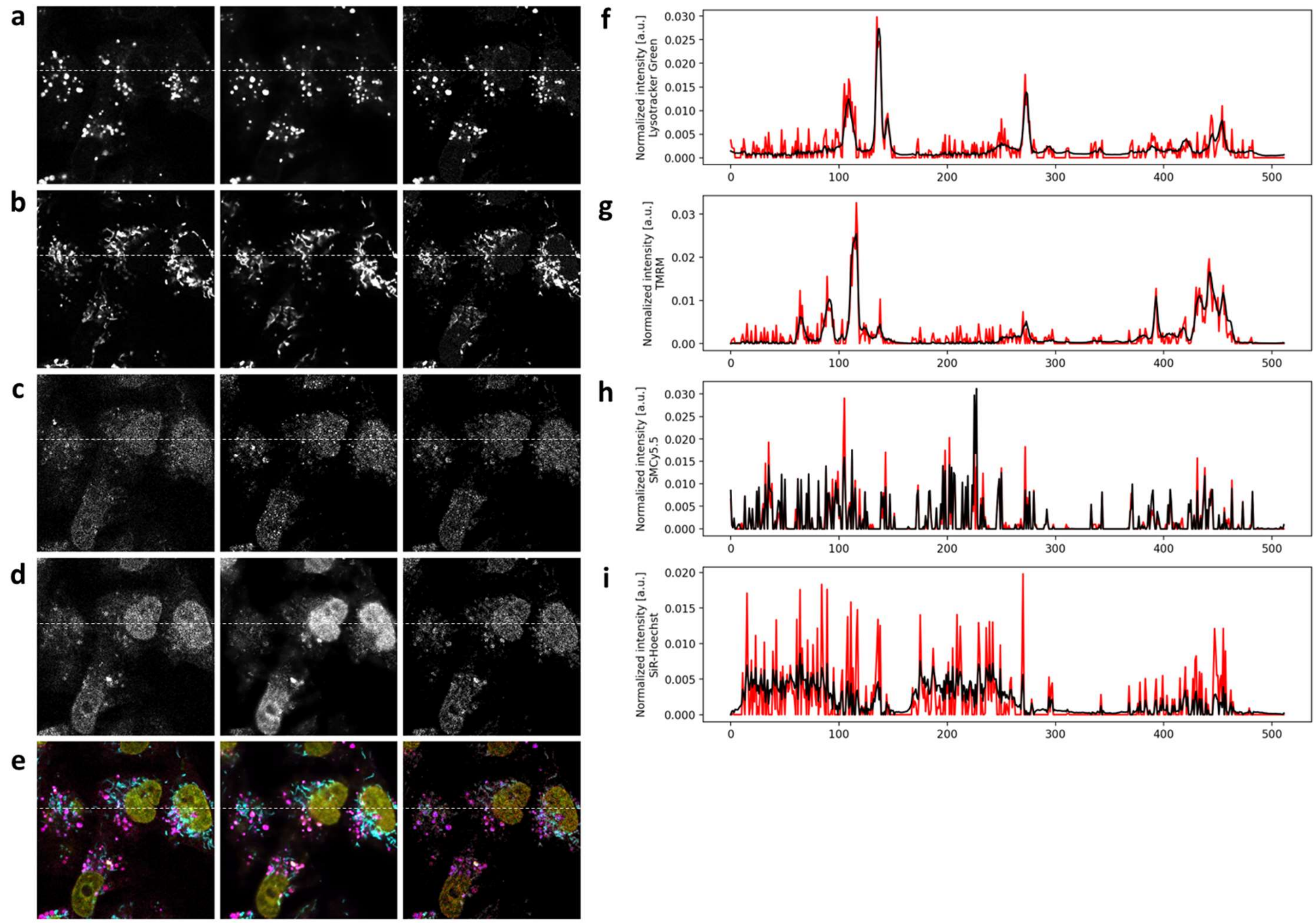


**_Figure S2: Denoising_**

*Single (**a-d**) and combined (**e**) channels of a representative image of MDA-MB-231 cells (magenta: lysosomes, cyan: mitochondria, green: lipid droplets, yellow: DNA). Single channels refer to lysosomes (**a**), mitochondria (**b**), lipid droplets (**c**) and DNA (**d**), respectively. Images are high-SNR (left), denoised (center) and low-SNR and graphs in **f-I** show the low-SNR (red line) and denoised (black line) along the white dashed line shown in **a-e.***

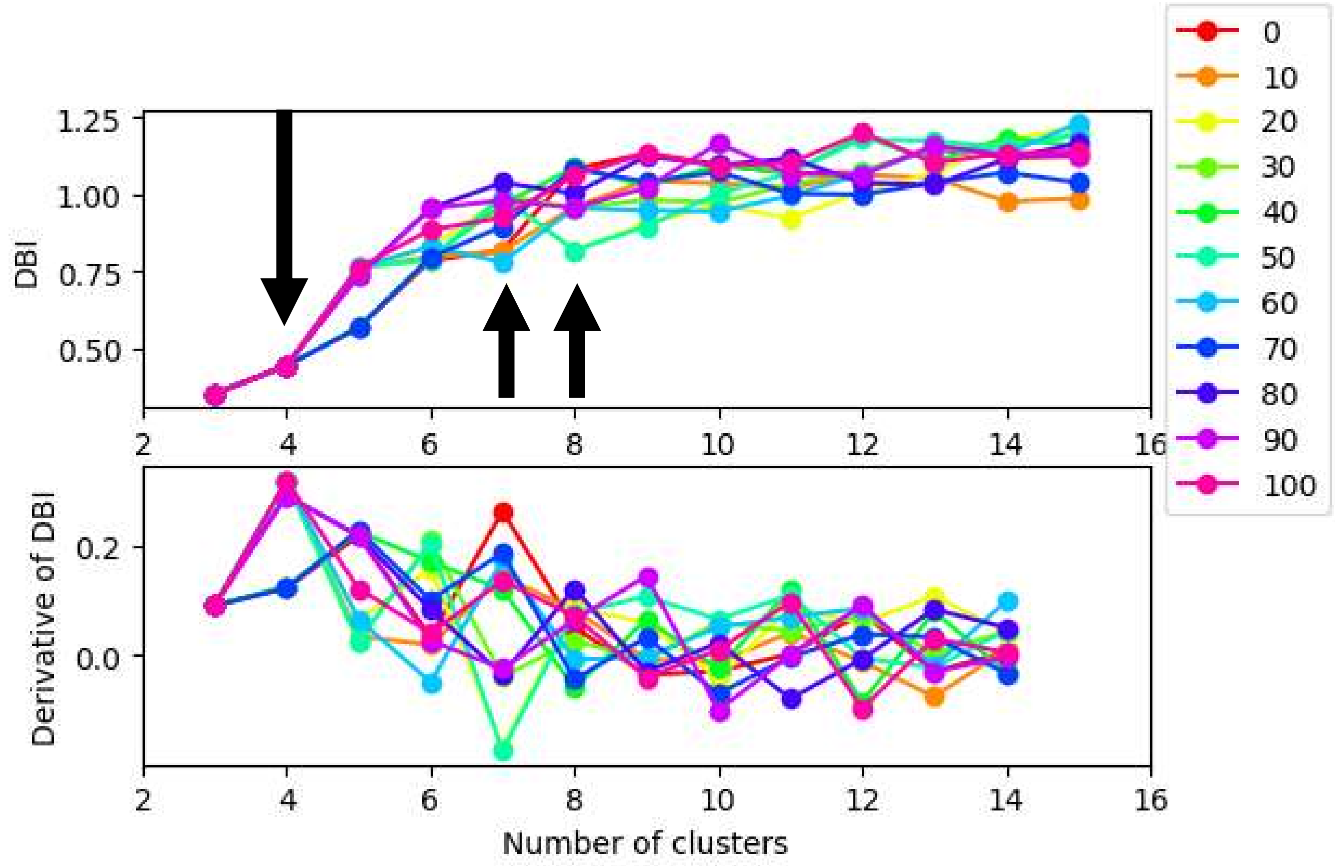


***<u>Figure S3: Clustering, number of clusters selection</u>***

*Davies-Bouldin Index (DBI) as a function of the number of clusters (top) used for GMM clustering for different starting conditions (0-100 in steps of 10) and its derivative (bottom). Black arrows highlight local minima in the DBI for specific numbers of clusters, corresponding to local maxima in the derivative.*

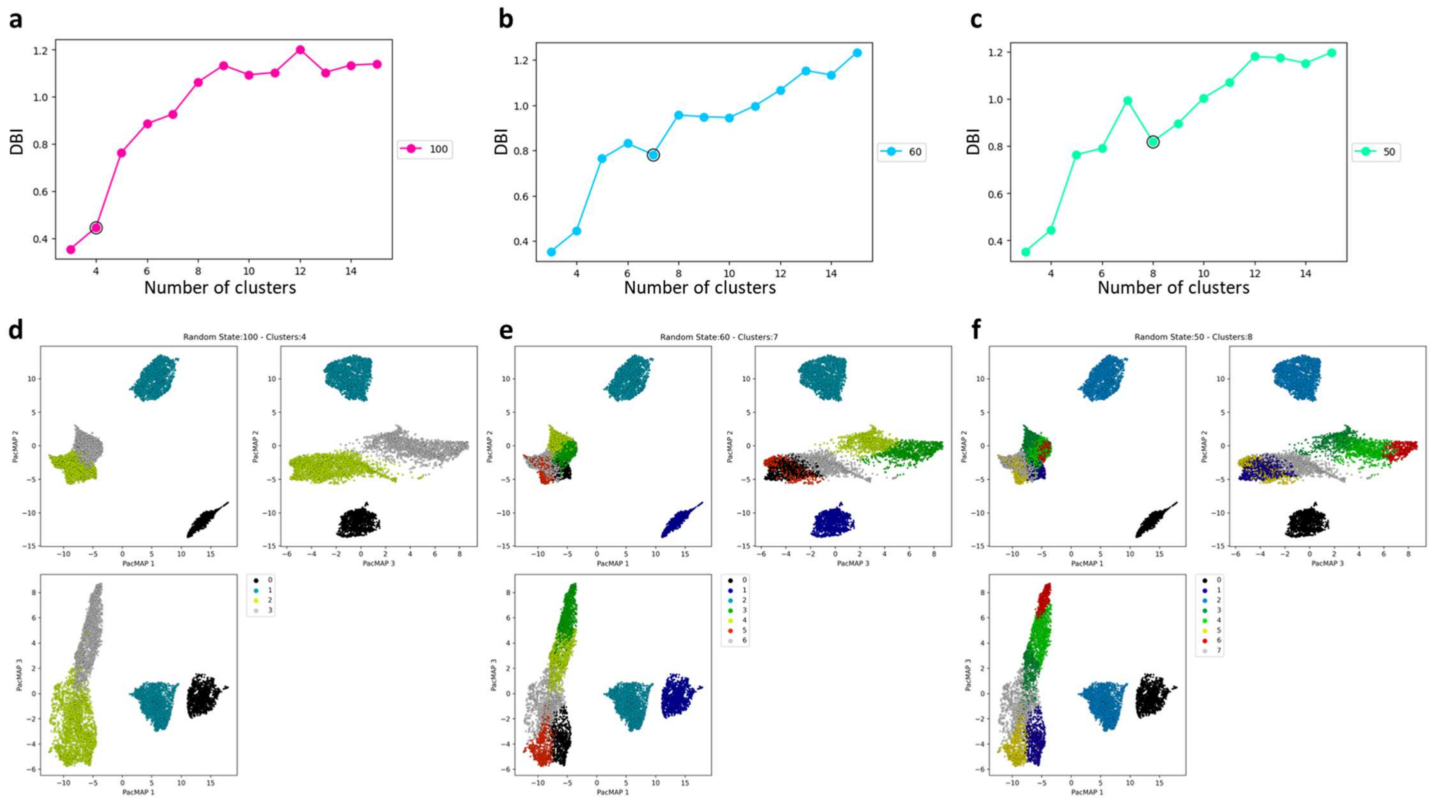


***<u>Figure S4: Clustering, example with different numbers of clusters</u>***

*Davies-Bouldin Index (DBI) as a function of the number of clusters (**a-c**) for the highlighted conditions in **Figure S3**, for different starting conditions (100, 60 and 50 for **a, b** and **c**, respectively), black circle highlights different possible numbers of clusters. PacMAP representation of ESPRESSO phenotypes from data in **Figure 1** with 4, 7 or 8 clusters (**d**, **e** and **f**, respectively), where different colors represent different clusters.*

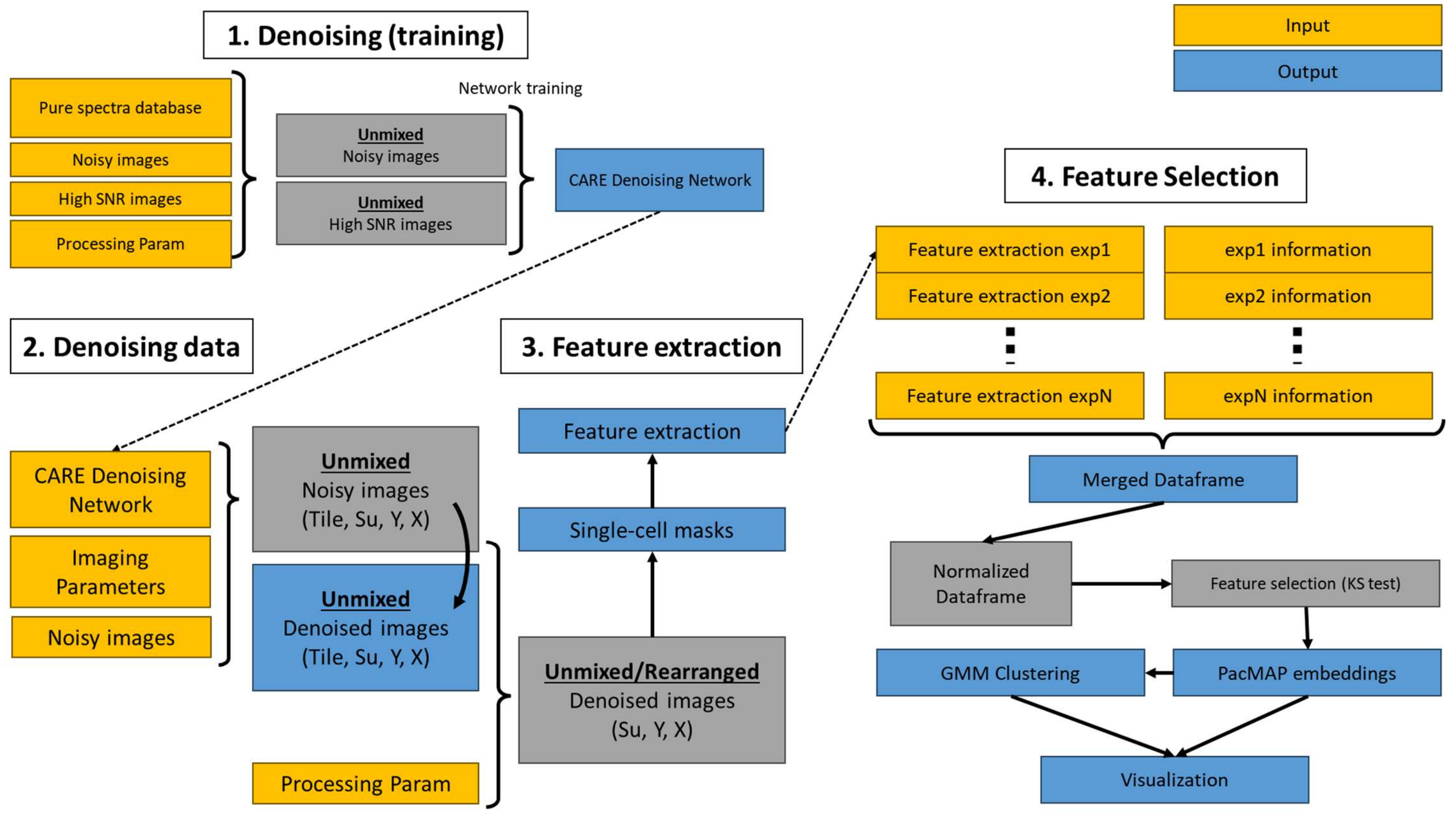


***Figure S5: Workflow***

*Schematic workflow of the ESPRESSO pipeline. First, a CARE denoising network is trained by inputting the pure spectra of the dyes used during imaging, a set of noisy images and a set of high SNR images, together with appropriate processing parameters. Both noisy and high SNR images are unmixed by spectral unmixing and used for training (see Methods), outputting a CARE denoising network for each of the unmixed channels. Second, a denoising step is applied to the experimental data, inputting noisy images and the appropriate denoising network and imaging parameters, returning unmixed, denoised image in the shape Tile (number of tiles) by Su (Spectrally unmixed channels) by Y by X (X and Y being spatial dimensions). Third, organelle features are extracted from the denoised images that are reconstructed (using the appropriate processing parameters) into denoised images in the shape of Su by Y by X, from which single-cell masks are calculated and the features are extracted (see Methods). Finally, data frames from multiple experiments are loaded and appropriately labeled, generating a merged data frame containing all cell features from all conditions considered. The data frame is normalized and the most relevant features are selected to construct PacMAP embeddings, from which GMM clusters are selected. Information from embeddings and clusters are used for visualization.*
